# A coelenterazine-dependent luciferase from the deep-sea coral *Anthoptilum murrayi* Kölliker, 1880 (Cnidaria: Octocorallia: Pennatulacea)

**DOI:** 10.64898/2025.12.27.696692

**Authors:** Gabriela A. Galeazzo, Douglas M. M. Soares, Danilo T. Amaral, Patricia Sartorelli, Ana Clara L. N. Silva, Eliana Samuels, Marcelo R. S. Melo, Cassius V. Stevani, Anderson G. Oliveira

## Abstract

Bioluminescence is the production of visible light by living organisms through biochemical reactions in which specialized enzymes known as luciferases catalyze the oxidation of light-emitting substrates (luciferins), producing photons. In the ocean, bioluminescence is widespread among anthozoans (corals and sea anemones), yet the molecular basis of their light emission remains poorly defined. To date, molecular characterization within Anthozoa has largely focused on the sea pansy *Renilla reniformis* (Octocorallia: Renillidae), leaving anthozoan luciferase diversity underexplored. Here, we report the identification and biochemical characterization of a coelenterazine-dependent luciferase from the deepsea sea pen *Anthoptilum murrayi* (AnmLuc). Transcriptome analysis identified a transcript encoding a ∼34-kDa protein bearing motifs characteristic of coelenterazine-dependent luciferases. The coding sequence was cloned and the recombinant protein was expressed in *Escherichia coli*. The purified enzyme produced intense blue emission (λ_max_ ≈ 495 nm) in the presence of coelenterazine and displayed an activity optimum near 5 °C. These findings provide the first molecular characterization of a luciferase from a deep-sea anthozoan, expand the known diversity of coelenterazine-dependent luciferases in Cnidaria, and offer new insights into the mechanism and evolution of light emission in Anthozoa.

## 1 Introduction

In bioluminescence, light emission results from enzyme-catalyzed oxidation in which a luciferase converts a small-molecule luciferin into an excited-state product that relaxes by emitting a photon [1]. Despite its broad occurrence across marine taxa, the biochemical basis of most bioluminescent systems remains unclear; only a minority have defined components, and fewer still have fully elucidated pathways [2]. In anthozoans, a group of cnidarians that includes sea anemones and corals, the canonical model is the sea pansy *Renilla reniformis* (Pallas, 1766) [3]. In this system, coelenterazine, an imidazopyrazinone-type luciferin found in both luminous and some non-luminous marine organisms, functions with a Ca^2+^-regulated luciferin-binding protein (LBP), a coelenterazine-dependent luciferase, and green fluorescent protein (GFP). An increase in Ca^2+^ triggers LBP to release coelenterazine, allowing luciferase to oxidize it to coelenteramide and CO2 with blue emission (λ_max_ ≈ 480 nm), and resonance energy transfer to GFP shifts the observed emission to green (λ_max_ ≈ 509 nm) [2].

Coelenterazine-dependent luciferase activity has been reported in several anthozoans of the superfamily Pennatuloidea, which includes both the sea pens (Pennatulacea) and the sea pansy (Renillidae), in which *Anthoptilum murrayi* Kölliker, 1880 and *Renilla reniformis* are found, respectively. Sea pens are key structural components of soft-bottom benthic ecosystems, and most species inhabit deep-sea environments (∼200–6000 m) [4]. Nevertheless, the biochemistry of light emission in deep-sea pennatulaceans remains largely unexplored. A recent work demonstrated the widespread occurrence of coelenterazine-dependent luciferases in deep-sea octocorals, including the families Isididae, Alcyoniidae, Umbellulidae, Funiculinidae, Kophobelemnidae and Protoptilidae [3]. Despite this distribution, the surface-dwelling sea pansy *Renilla reniformis* remains the only anthozoan for which the complete bioluminescent system has been isolated and cloned [2]. Identifying and characterizing additional luciferases in these taxa is essential for clarifying the evolution of bioluminescent systems and for discovering enzymes with biotechnological applications.

Here, we report the biochemical characterization of a novel coelenterazine-dependent luciferase (AnmLuc) from the deep-sea coral *Anthoptilum murrayi*. Colonies from the South Atlantic were collected during our deep-sea research surveys. *A. murrayi* produces green light when mechanically stimulated (λ_max_ ≈ 515 nm). The recombinant enzyme emits bright blue light upon addition of coelenterazine (λ_max_ ≈ 495 nm) and displays a similar substrate affinity to RLuc (K_M_ = 3.39 µM vs 2.9 µM for RLuc) [5]. Unlike the *Renilla* luciferase, AnmLuc is cold-adapted, with maximal activity near 5°C, reflecting its deep-sea origin [6]. These results constitute the first functional characterization of a luciferase from a deep-sea anthozoan and highlight its utility for sensitive bioluminescent assays under low-temperature conditions.

## 2. Material and Methods

### 2.1 Collection of A. murrayi

Specimens of the deep-sea coral *A. murrayi* were collected during the DEEP-OCEAN expedition (September 2019) on the Brazilian continental slope off São Paulo (24°01′–24°50′ S, 43°48′–45°51′ W) aboard R/V *Alpha Crucis*. Sampling was conducted at 200–1,500 m depth (captures at ∼1,100 m) using a demersal otter trawl (17 m groundrope, 2.5 m vertical opening; 10 cm mesh in wings/body and 2.5 cm in the codend). Net-sensor oceanographic data were recorded with Marport instrumentation. Specimens were identified onboard, flash-frozen in liquid nitrogen, and stored at −80 °C until use. Collections were authorized by SISBIO (permits 28054-4 and 82624-1) and the Brazilian Navy’s Interministerial Commission for Marine Resources (Ordinance No. 223).

### 2.2 UHPLC–ESI–HRMS/MS Analysis of Methanolic Extracts of *A. murrayi*

A methanolic extract was prepared by homogenizing ∼8 mg frozen coral tissue in 1 mL methanol using an Ultra-Turrax homogenizer (IKA T10 basic). Homogenates were centrifuged (20,000 × *g*, 10 min, 4 °C), and 5 µL of the supernatant was injected for LC–MS/MS analysis. Separations were performed on a Shimadzu Prominence system equipped with a diode-array detector (SPD-M20A) using a C18 reversed-phase column (Kinetex, 2.6 µm, 100 × 2.1 mm) at 30 °C and a flow rate of 0.2 mL min^-1^. Mobile phases were (A) 0.1% (v/v) formic acid in water and (B) acetonitrile, with the following gradient: 5% B (0–2 min), 5–100% B (2–10 min), 100% B (10–14 min), 100–5% B (14–15 min), and 5% B (15–18 min). The column effluent was analyzed on a Bruker microTOF-QII QTOF mass spectrometer operating in positive ESI mode (*m/z* 100–1200; ∼18,000 FWHM). Source conditions were: 200 °C, N2 drying gas 5 L min^-1^, capillary 4000 V, and cone 500 V. Mass calibration was performed with sodium formate introduced at 10 µL min^−1^ via syringe pump (KD Scientific).

### 2.3 Identification of luciferase-like sequences in *A. murrayi*

The *de novo* transcriptome of *A. murrayi* employed in this study was sequenced and assembled as previously described by Duchatelet et al. [7]. Briefly, total RNA was extracted from whole organism tissue, and the cDNA library was sequenced using the Illumina HiSeq2500 platform (150 bp paired-end reads). The raw reads were filtered for quality control and assembled using Trinity v2.13.2. Transcriptome completeness was assessed using BUSCO v4 (Metazoa dataset), revealing 94.4% completeness. Local tBLASTn searches of this transcriptome were then performed using *Renilla* luciferase (RLuc) as the query, as previously described [7]. Putative unigenes were validated by reciprocal BLASTx against the NCBI non-redundant database. Open reading frames (ORFs) were identified using ExPASy Translate (https://web.expasy.org/translate/), and predicted proteins were aligned with homologs from other metazoans in Geneious for motif annotation. Conserved features of *Renilla*-type luciferases, including the catalytic triad, were verified [7]. Among candidates, Comp24811 (AnmLuc) was the most highly expressed and satisfied all criteria (single high-confidence hit, complete ORF, conserved catalytic motifs) [8,9]; we therefore selected it for cloning and expression.

### 2.4 Cloning, expression and purification of recombinant *A. murrayi* luciferase (AnmLuc)

The AnmLuc coding sequence was synthesized with *E. coli* codon optimization and cloned into pET-28a(+) (Biomatik) to generate an N-terminal His6 fusion (BamHI/HindIII). The pET-28a(+)-AnmLuc plasmid was transformed into *E. coli* ArcticExpress (DE3) (Agilent) by heat shock (42 °C, 2 min) and recovered in LB (37 °C, 1 h). Transformants were grown in LB + kanamycin (50 µg/mL) at 11 °C (180 rpm, 18 h) and induced with IPTG (1 mM, 2 h, 11 °C). Cells were harvested and resuspended in lysis buffer (50 mM sodium phosphate pH 7.4, 300 mM NaCl, 10 mM imidazole, 1 mM PMSF, 2 mM β-mercaptoethanol) with lysozyme (1 mg/mL), disrupted on ice with a Turrax homogenizer (15 × 10 s pulses), and clarified by centrifugation (10,000 × *g*, 30 min, 4 °C). The soluble fraction was incubated with Ni–NTA resin (1 h, 4 °C), loaded onto a gravity column, washed (3 × 10 mL lysis buffer), and eluted with an imidazole step gradient (40 mM then 200 mM; 5 mL each). Fractions were analyzed by SDS– PAGE, pooled, and buffer-exchanged into 50 mM Tris–HCl (pH 7.5), 70 mM NaCl, and 1 mM DTT.

### 2.5 Light emission assays

Assays were performed on a Sirius L single-tube luminometer (Berthold). Reactions (500 µL) contained recombinant luciferase (100 µL; 150–450 µg mL^-1^) in 50 mM sodium phosphate (pH 7.4) with 70 mM NaCl. Reactions were initiated by adding native coelenterazine (NanoLight/Prolume, Pinetop, AZ, USA) to 1.5 µM unless otherwise noted. Coelenterazine concentration was determined by absorbance at 435 nm (ε435 = 9,800 M^-1^·cm^-1^) [1]. Luminescence was recorded for 60 s at 25 °C (triplicate). For pH profiling, 50 mM buffers spanning pH 4.0–10.0 were used (sodium citrate, pH 4–5; sodium phosphate, pH 6–7; Tris–HCl, pH 8–10). Ionic strength was varied by supplementing the assay buffer (50 mM sodium phosphate, pH 7.4) with NaCl (30–500 mM). Temperature dependence/thermostability was assessed from 5–40 °C using a ThermoMixer (Eppendorf); enzyme aliquots were pre-incubated for 10 min at each temperature and measured immediately in the luminometer chamber at the same temperature. Kinetic measurements (RLU/s) were obtained at 25 °C in 50 mM sodium phosphate (pH 7.5) across the indicated coelenterazine concentrations (n = 3 per concentration). K_m_ and V_max_ were obtained by nonlinear regression in GraphPad Prism 10.6.1 using a Michaelis–Menten model, fitting the mean rate at each concentration with 1/SD^2^ weighting (SD from the triplicates). Parameter uncertainty was estimated using profile-likelihood 95% confidence intervals.

### 2.6 Spectral measurements

Frozen *A. murrayi* tissues were thawed, rehydrated in deionized water, and bioluminescence spectra were recorded at room temperature using a cooled CCD spectro-imaging system (LumiF SpectroCapture AB-1850). Spectra were collected with 2-min exposures (n = 3). For in vitro spectra, 100 µL of AnmLuc was mixed with 397 µL of 50 mM sodium phosphate (pH 7.4) containing 70 mM NaCl, and the reaction was initiated by adding 3 µL of 1 mM coelenterazine (final volume, 500 µL; final [coelenterazine], 6 µM).

### 2.7 AlphaFold2 structure prediction

Protein structures were predicted with AlphaFold2 via ColabFold [8,9]. The AnmLuc model was compared with *Renilla* luciferase (RLuc; PDB 7OMR), and GFP-related predictions were generated for *Renilla* GFP (RGFP; PDB 2RH7) and an AnmGFP Y66 variant (comp24361_c0_seq1 with Tyr restored at the putative fluorophore-forming position). Top-ranked models were used for analyses; confidence was assessed by per-residue pLDDT. Visualization and superposition were performed in UCSF ChimeraX [10]

## 3. Results

### 3.1 Coelenterazine drives luciferase activity in *A. murrayi*

Homogenates of *A. murrayi* tissue (Tris–HCl, 50 mM, pH 7.4) emitted detectable light only after the addition of native coelenterazine, confirming strict substrate dependence. Upon substrate addition, the signal increased by approximately six orders of magnitude above baseline. Methanolic tissue extracts analyzed by UHPLC–ESI–HRMS/MS identified coelenterazine and its oxidation product coelenteramide, as well as hydrolysis products, based on retention times and *m/z* values (Supplementary Fig. S1).

### 3.2 Transcriptome-guided identification and production of recombinant *A. murrayi* luciferase

Assembly and annotation of transcriptomic data from luminous *A. murrayi* tissue [7] identified a candidate luciferase transcript (comp24811_c0_seq1) encoding a 304-aa protein (predicted 34.6 kDa) with conserved catalytic triad/active-site residues characteristic of coelenterazine-dependent luciferases. BLAST comparison to RLuc (311 aa) showed 58% identity with 98% query coverage [11] (Fig. 1). To test function, the coding sequence was cloned into pET-28a(+) and expressed in *E. coli*. Expression in BL21(DE3)-RIL yielded predominantly insoluble protein across induction conditions, and post-lysis solubilization (2–8 M urea, 2% Triton X-100, 1% CHAPS) was unsuccessful. Soluble recombinant AnmLuc was obtained only in *E. coli* ArcticExpress (DE3) induced overnight at 11 °C with enhanced lysozyme treatment. Ni–NTA purification yielded a dominant ∼35 kDa band eluting at 200 mM imidazole (with nonspecific bands eluting at 40–100 mM; Supplementary Fig. S2), and the eluate emitted light upon coelenterazine addition in phosphate buffer (50 mM, pH 7.4), confirming recovery of an active recombinant luciferase.

**Fig. 1:**
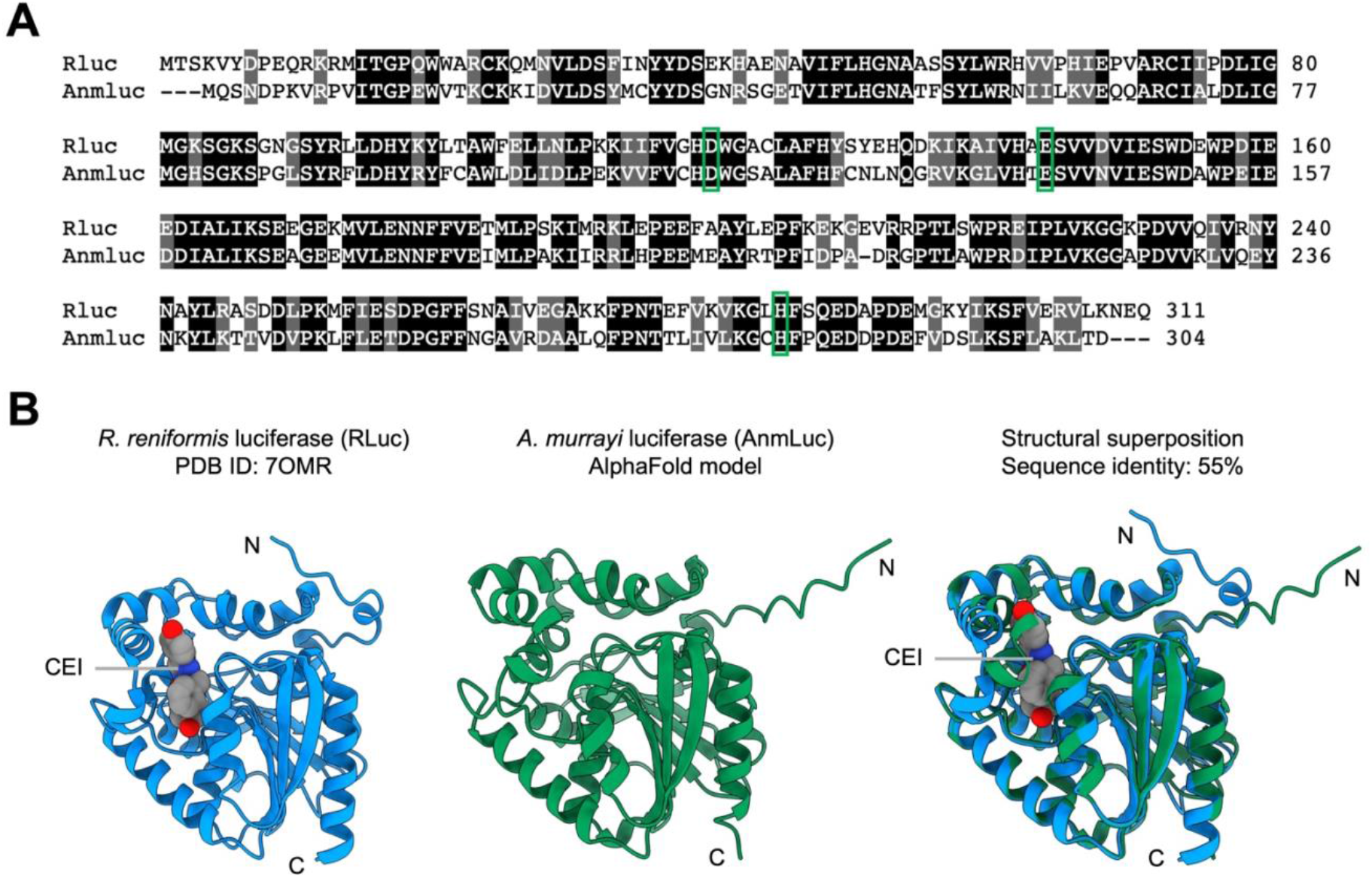
(A) Multiple sequence alignment of *R. reniformis* luciferase (RLuc) and *A. murrayi* luciferase (AnmLuc). Alignment was generated with Clustal Omega and visualized with Color Align Conservation; identical residues are shown in black and similar residues in gray. Residues corresponding to the conserved catalytic triad are boxed in green. (B) Structural comparison of RLuc (PDB 7OMR; bound CEI, coelenteramide) and the AlphaFold2 model of AnmLuc. Structures are shown as cartoons (RLuc, blue; AnmLuc, green) with N- and C-termini indicated. Right panel shows the structural superposition of AnmLuc onto RLuc (pairwise sequence identity ∼55%).

### 3.3 Characterization of *A. murrayi* recombinant luciferase

Light-emission assays confirmed that the purified fraction contained an active luciferase corresponding to a ∼35 kDa band, and this material was used for all subsequent characterization (Fig. 2A). AnmLuc showed an optimal reaction pH of 7.0 (Fig. 2B), similar to reported RLuc behavior [1,12–14]. Ionic strength profiling (30–500 mM NaCl) indicated maximal activity at ∼70 mM NaCl (Fig. 2C). Coelenterazine titrations (0.04–50 µM) yielded K_M_ = 3.39 µM and V_max_ = 1.09 × 10^5^ RLU/s, (R^2^ = 0.942) (Supplementary Fig. S3). Temperature profiling indicated a psychrophilic enzyme with maximal activity near 5 °C (Fig. 2D), consistent with its deep-sea origin and distinct from RLuc (25 °C) [6,15,16].

**Fig. 2:**
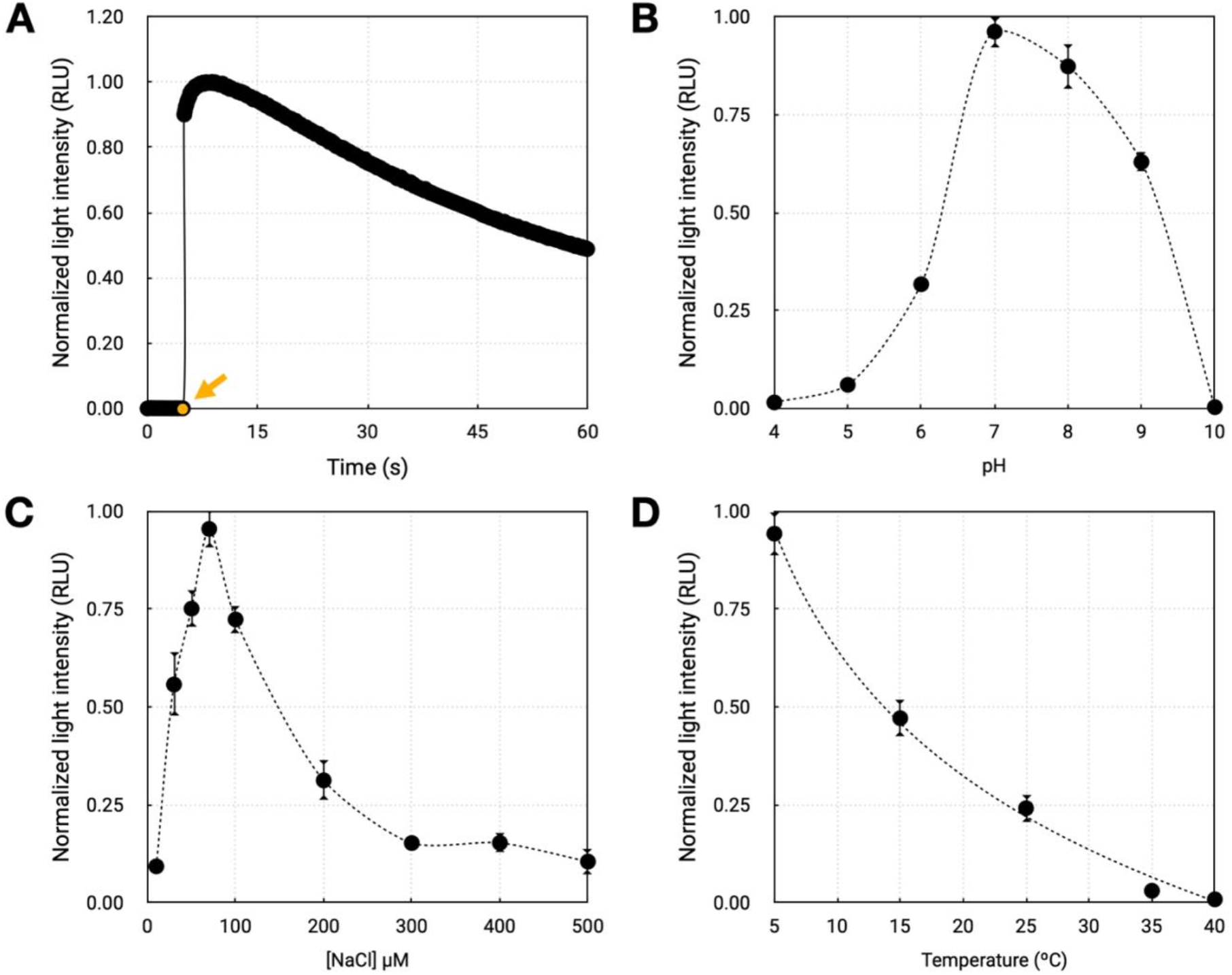
(A) Luminescence time course of purified AnmLuc; the arrow indicates coelenterazine addition to initiate the reaction. (B) pH dependence of AnmLuc luminescence (pH 4–10). (C) Effect of ionic strength on AnmLuc luminescence across NaCl concentrations (30–500 mM). (D) Temperature dependence of AnmLuc luminescence (5–40 °C).

### 3.4 Emission Spectrum Analysis

The emission spectrum of the purified recombinant AnmLuc was measured, revealing a blue bioluminescence with a maximum emission at 495 nm (Fig. 3A). This *in vitro* blue emission contrasts with the green light (λ_max_ ≈ 515 nm) observed *in vivo* from the coral tissue (Fig. 3B). This spectral shift strongly suggests that the *in vivo* green color is produced by Bioluminescence Resonance Energy Transfer (BRET) from the blue-emitting luciferase to an accessory fluorescent protein, which was absent in the recombinant assay [17–19].

**Fig. 3:**
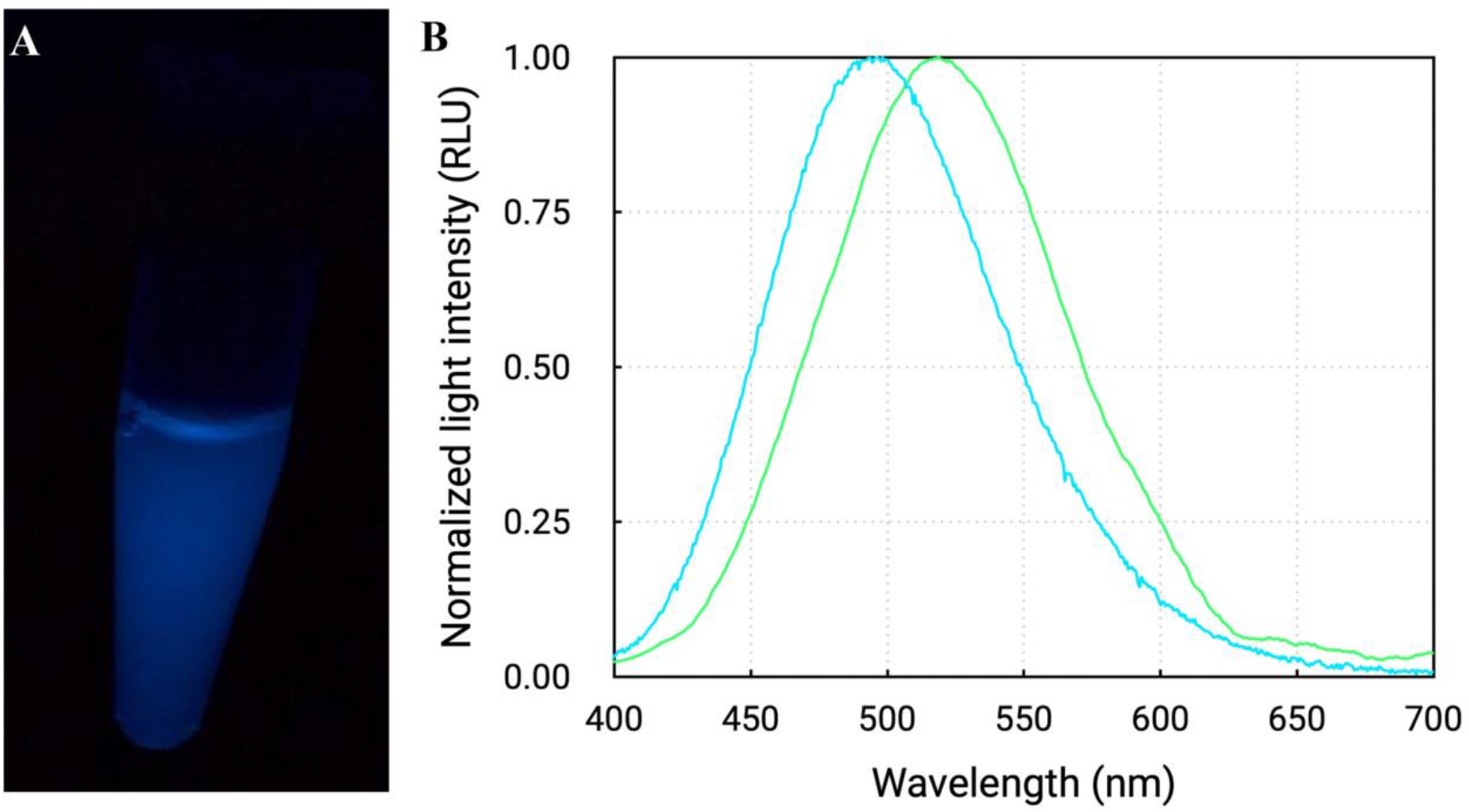
(A) Photograph of AnmLuc emitting blue light following addition of coelenterazine. (B) Emission spectra of AnmLuc (blue line) and coral tissue (green line).

**Fig. 4:**
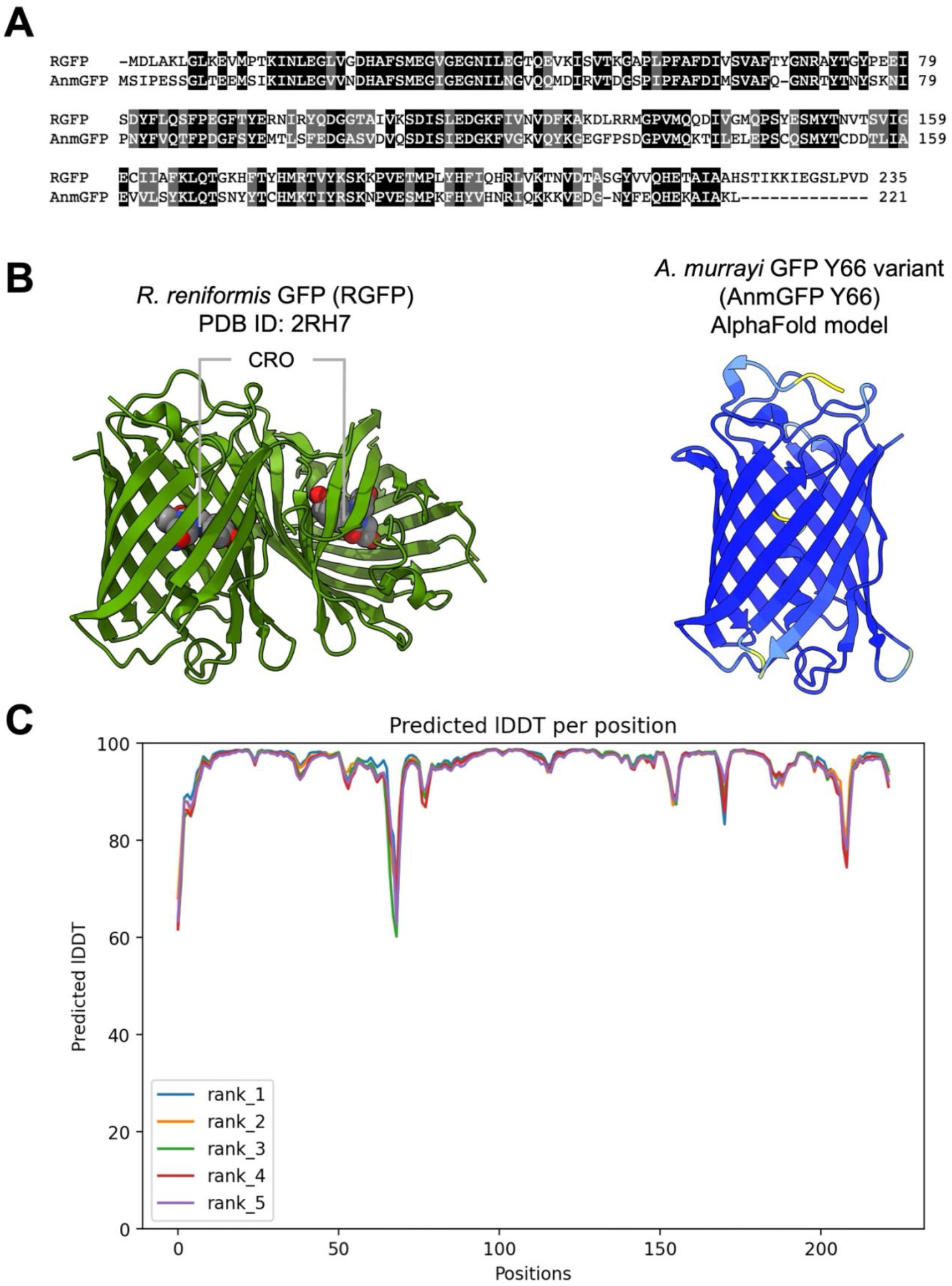
(A) Multiple sequence alignment of *R. reniformis* GFP (RGFP) and *A. murrayi* GFP (AnmGFP). Alignment was generated with Clustal Omega and visualized with Color Align Conservation; identical residues are shown in black and similar residues in gray. (B) RGFP crystal structure (PDB: 2RH7; dimer); chromophore (CRO) indicated. (C) AlphaFold2 model of AnmGFP Y66 (cartoon) and predicted confidence (pLDDT; rank_1–rank_5) plotted as predicted lDDT per position.

## 4. Discussion

The bioluminescent system of *Anthoptilum murrayi* relies on coelenterazine, as in other octocorals [1,15]. Addition of native coelenterazine to coral protein extracts triggered strong light emission, indicating strict substrate dependence. LC–MS/MS of methanolic tissue extracts detected endogenous coelenterazine and its oxidation product coelenteramide (Supplementary Fig. S1), establishing coelenterazine as the native luciferin of *A. murrayi*, analogous to *R. reniformis* [20]. Transcriptome analysis identified the main molecular components of the bioluminescent system, including a highly expressed luciferase candidate (AnmLuc) and several GFP-like sequences. Recombinant AnmLuc expression poses solubility challenges typical of coelenterazine-dependent luciferases in prokaryotic hosts [2]. Although expression in *E. coli* BL21(DE3)-RIL was successful, most of the protein accumulated in the insoluble fraction. Multiple approaches (urea solubilization, SUMO fusion, and expression in *Saccharomyces cerevisiae*) failed to yield active enzyme. Soluble, active AnmLuc was obtained only in *E. coli* ArcticExpress®, which coexpresses cold-adapted chaperonins (Cpn60/Cpn10); induction at 11 °C and optimized lysis enabled proper folding and recovery of the recombinant luciferase.

Functional characterization of the enzyme revealed a typical “flash”-type luminescence profile (Fig. 2A), with a rapid emission peak followed by a sharp decay, similar to RLuc [21]. AnmLuc exhibited maximal activity at pH 7.0, consistent with a neutrophilic profile (Fig. 2B). Its optimal temperature of 5 °C indicates a psychrophilic nature, consistent with the deep-sea habitat of *A. murrayi* (Fig. 2D). Maximum activity was observed at 70 mM NaCl, decreasing at higher salt concentrations, suggesting sensitivity to ionic strength (Fig. 2C). Michaelis–Menten kinetic analysis revealed a K_M_ of 3.39 µM like those reported for RLuc, whose K_M_ values typically range from 2 to 5 µM. The measured V_max_ was 1.09 × 10^5^ RLU s^−1^, while the native RLuc has been reported to exhibit a turnover number corresponding to a V_max_ of approximately 1.85 s^−1^ per mol of enzyme under substrate-saturating conditions [1,22]. While the similar K_M_ indicates comparable substrate affinity, the lower V_max_ reflects a reduced maximal catalytic activity relative to RLuc under our conditions, consistent with differences in steps beyond substrate binding. Nevertheless, the K_M_ in the low-µM range supports robust activity at modest coelenterazine concentrations and suggests potential utility of AnmLuc as a biotechnological reporter (Supplementary Fig. S3).

AnmLuc consists of 304 amino acids and has a molecular mass of 34.6 kDa, values similar to those of RLuc, which contains 311 residues and approximately 37 kDa [21]. Sequence alignment revealed 58% identity and 98% coverage between the two enzymes, indicating strong structural conservation. Predictive modeling based on AlphaFold (Jumper et al., 2021) showed that AnmLuc retains the typical α/β-hydrolase fold with a cap-domain and a conserved catalytic triad (D120, E144, H285), as described for RLuc [8]. Residues implicated in oxygen binding and stabilization during the bioluminescent reaction, including N53 and W121, were also conserved in AnmLuc, further supporting the maintenance of the canonical RLuc catalytic mechanism [7].

Interestingly, some substitutions observed in AnmLuc occur at positions analogous to mutations artificially introduced in RLuc8, an engineered variant designed for enhanced stability and catalytic efficiency [5]. From an evolutionary perspective, RLuc-type luciferases are homologous to bacterial haloalkane dehalogenases, suggesting that cnidarian luciferases may have originated from an ancient horizontal gene transfer event [7]. Furthermore, the presence of a coelenterazine-dependent RLuc-based system across multiple pennatulacean families, including *A. murrayi* and previously described shallow-water species, indicates that this bioluminescent system arose prior to the diversification of Pennatulacea and has been vertically inherited with strong functional conservation [7].

Finally, a notable discrepancy emerged between the *in vivo* and *in vitro* emission spectra. While coral tissue emits green light (∼515 nm), the purified recombinant luciferase emits blue light (∼495 nm) (Fig. 3). In bioluminescent systems such as *R. reniformis*, this blue-to-green spectral shift can arise from BRET between the luciferase and an associated GFP [17,23]. Transcriptomic analysis of *A. murrayi* identified several GFP-like sequences; however, the most highly expressed candidate (comp24361_c0_seq1) lacks the conserved Y required for maturation of the canonical chromophore (SYG), with the Y absent from the corresponding QYG region [19,24]. In addition, other candidate sequences exhibit an altered N-terminus and a truncated C-terminus, lacking the extended C-terminal region implicated in oligomerization/association in some cnidarian GFPs. Consistent with these features, preliminary assays detected no fluorescence upon recombinant expression, even after restoring Y at the putative chromophore position (Y66) by site-directed mutagenesis.

AlphaFold2 predictions for *Renilla* GFP (RGFP; the canonical GFP reference) and AnmGFP Y66 supported a conserved GFP-like β-barrel core but indicated reduced confidence in specific loop and terminal regions of the *A. murrayi* sequences; importantly, Y66 restoration did not substantially alter the predicted fold or confidence profile (Supplementary Fig. S3). Together, these observations suggest that the green emission observed in coral tissue may reflect a non-uniform (region-specific) abundance and/or association of GFP-like proteins with luciferase across the colony. This interpretation is consistent with prior reports in the sea pen *Umbellula magniflora*, where emission collected from different body regions shifts from a broader blue signal near ∼470 nm to a narrower, longer-wavelength band near ∼500 nm, consistent with variable GFP coupling along the rachis: spectra at the top resemble the in vitro, uncoupled (blue) emission, whereas spectra nearer the peduncle show GFP-shifted emission [25–27]. Additional experiments are underway to test this model, including region-resolved spectral measurements across the colony and localization/quantification of GFP-like proteins relative to luciferase. Together, our results broaden the known diversity of coelenterazine-dependent systems and highlight deep-sea sea pens as a source of bioluminescent mechanisms with evolutionary and biotechnological relevance.

## CRediT authorship contribution statement

**Gabriela A. Galeazzo:** Validation, Methodology, Investigation, Writing - Original Draft, Visualization. **Douglas M. M. Soares:** Formal analysis, Writing - Review & Editing. **Danilo T. Amaral:** Formal analysis, Resources, Writing - Review & Editing. **Patricia Sartorelli:** Formal analysis, Resources, Writing - Review & Editing. **Ana Clara L. N. Silva:** Formal analysis, Writing - Review & Editing. **Eliana Samuels:** Formal analysis, Writing - Review & Editing. **Marcelo R. S. Melo:** Resources, Writing - Review & Editing. **Cassius V. Stevani:** Resources, Writing - Review & Editing. **Anderson G. Oliveira:** Supervision, Writing - Review & Editing, Funding acquisition, Project administration.

## Acknowledgements

We thank Prof. Vadim Viviani (Federal University of São Carlos, UFSCar) for access to a high-sensitivity spectroluminometer and Mr. Gabriel Pelentir for technical assistance.

## Supplemental data captions

**Supplementary Fig. S1**. Representative UHPLC–ESI–HRMS/MS chromatograms of the methanolic extract from coral tissue. (A) Base peak chromatogram of the methanolic extract from coral, in which peak 3 corresponds to coelenteramide and peak 5 to coelenterazine; (B) Extracted ion chromatograms (EICs) for coelenterazine (*m/z* 301.1; Rt 10 min); (C) EIC for coelenteramide (m/z 412.1; Rt 7.4 min).

**Supplementary Fig. S2**. Proteins were resolved by SDS–PAGE on a 12% gel and stained with Coomassie Brilliant Blue and molecular weight marker (kDa). Lane 1, Ni–NTA flow-through; Lane 2, wash; Lane 3, elution (40 mM imidazole); Lane 4, elution (60 mM imidazole); Lane 5, elution (80 mM imidazole); Lane 6, elution (100 mM imidazole); Lane 7, elution (200 mM imidazole). The expected AnmLuc band is indicated (∼35 kDa).

**Supplementary Fig. S3**. AnmLuc light emission (RLU/s) as a function of coelenterazine concentration. Points show mean ± SD (n = 3). The curve is a Michaelis–Menten fit in GraphPad Prism using 1/SD^2^ weighting (K_m_ = 3.39 µM; V_max_ = 1.09 × 10^5^ RLU/s; R^2^ = 0.942).

