## Supplementary figures and images for "A coelenterazine-dependent luciferase from the deep-sea coral *Anthoptilum murrayi* Kölliker, 1880 (Cnidaria: Octocorallia: Pennatulacea)"

### Supplementary Fig. S1

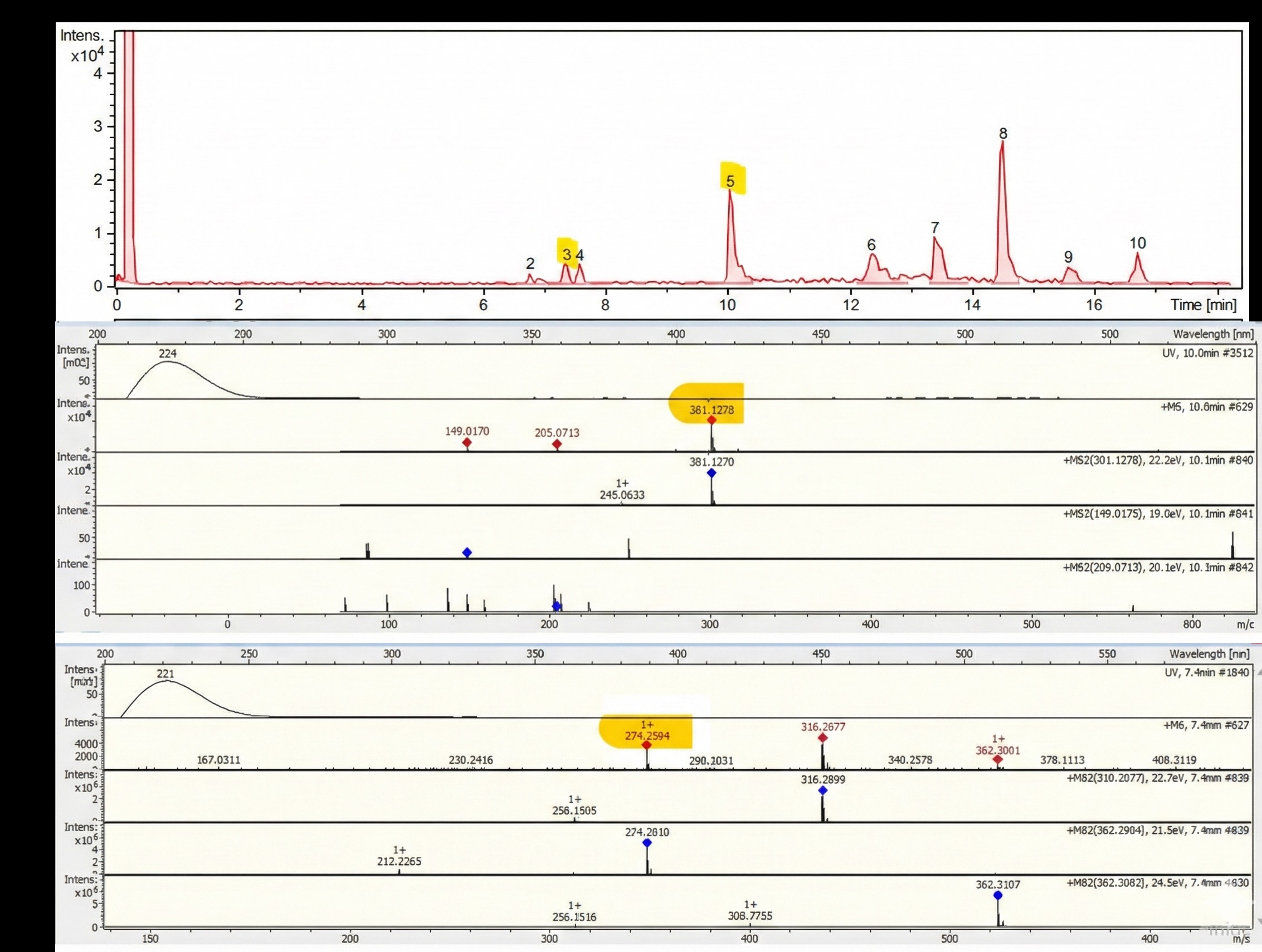

### Supplementary Fig. S2

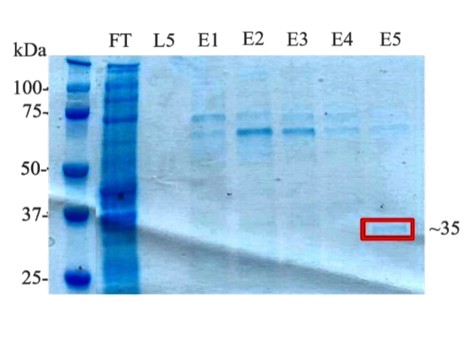

### Supplementary Fig. S3

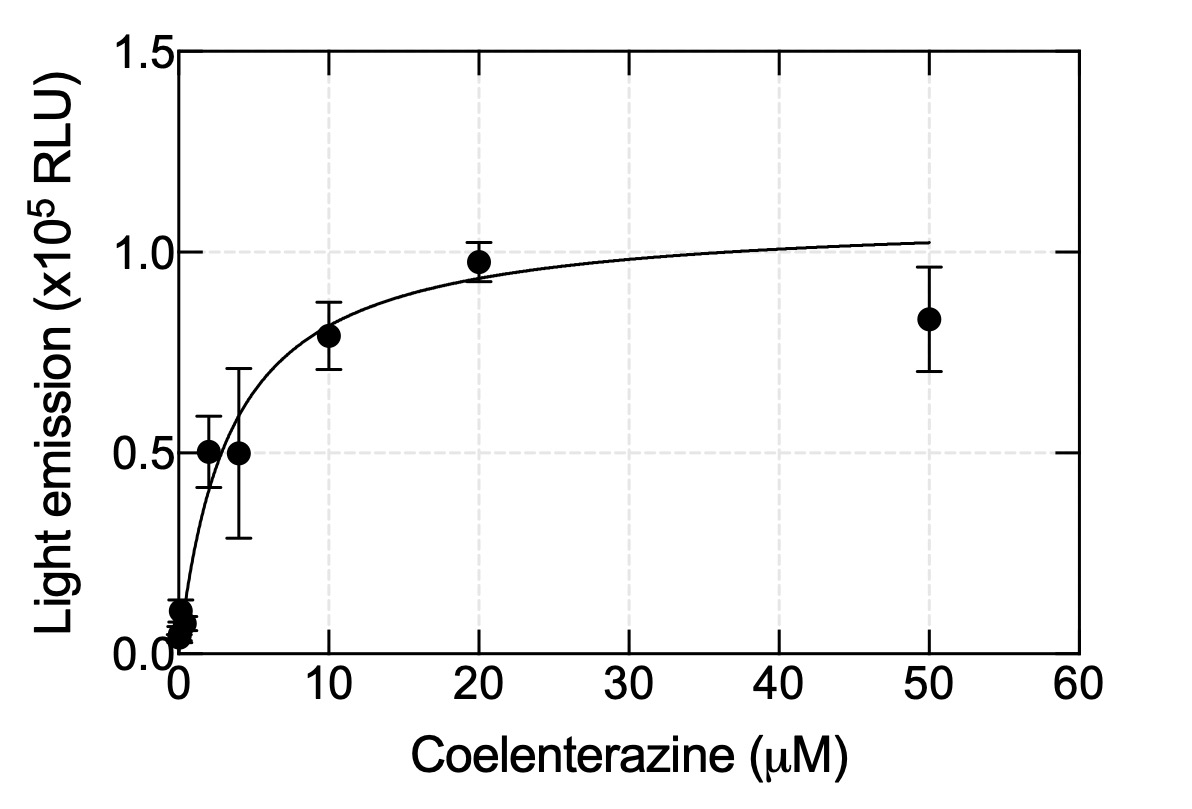
